# The Neural Protective Effects of Intermittent Fasting in Adult *Drosophila* Following Traumatic Brain Injury Exposure

**DOI:** 10.64898/2026.10.12.758363

**Authors:** Jesse J. Rojas, Marcus Intal, Andrew Kawwa, Celine Thomassian, Robert Squier, Sydney Mudrak, Kayla Lawani, Kim D. Finley

## Abstract

Traumatic brain injury (TBI) remains a major cause of neurological-based morbidity and mortality in different human populations. Repetitive, mild traumatic brain injury (**mTBI**) is increasingly recognized as a contributor to progressive neural attrition. Dietary measures that are neural protective could provide an accessible means for impacted human populations to alter outcomes in relation to neurological diseases. Intermittent fasting (**IF**) has been associated with metabolic and neuroprotective benefits for several neurological disease models, including *Drosophila melanogaster* models that are used to study neurological disorders linked to TBI exposure. Using two IF regimens, we examined the outcomes of adult *Drosophila* in conjunction with aging and mTBI. Overall, when exposed to trauma, IF-treated adult cohorts exhibited longer lifespans, lower mortality rates and improved behavioral responses. This included young (10-day) and middle aged (21-day) flies exposed to either Pre- or Post-IF conditions. Additionally, expression analysis showed IF treatments significantly reduced expression levels of *Drosophila* inflammatory markers (AMPs) in adult neural tissues from traumatized flies. The indications are that the net positive impact of IF found for longevity and trauma responses could be due to lower levels of secondary neuronal damage.

## Introduction

Traumatic Brain Injury (**TBI**) represents a substantial global health burden and a primary cause of chronic neurological disability and mortality across a wide range of populations. In the United States, foundational surveillance reports indicate that approximately 2.8 million TBI-related emergency department visits, hospitalizations, and deaths occur each year.^1^ In 2020, the Centers for Disease Control and Prevention (**CDC**) estimates there were 214,110 TBI related hospitalizations in the United States.^2^ By 2023, approximately 68,663 TBI-related deaths were documented, which doesn’t consider more minor injuries that resulted in emergency room, primary care, or urgent care visits. Further, CDC reports indicate that people, 75-years and older account for a higher percentage of TBI-related hospitalizations (32%) and deaths (28%).^3^ Overall, the rate of TBI-related Emergency Department (ED) visits result from common falls (413.2 per 100,000), being struck by or against an object (142.1) or are due to motor vehicle accidents (121.7).^2^

Repetitive, mild traumatic brain injury (**mTBI**) is now recognized as a chronic, progressive neurodegeneration and neural attrition over time.^4^ Generally, mTBI is classified as a concussion in which patients sustain alteration of consciousness no longer than 24 hours.^5,6^ Patients exposed to repetitive mTBI can exhibited many of the same negative outcomes and pathological neurodegenerative markers seen with a single, severe TBI event.^4^ Despite these consequences, a survey amongst high school football players found that only 47.3% young people went on to report their concussions, highlighting a clear understanding disconnect with reporting potentially life changing injuries.^7^ TBI underreporting is further exacerbated with domestic and other patterns of interpersonal violence, further delaying access to care.^8,9^ Overall, the pattern is that mild traumatic brain injury is widespread and underreported, which is compounded if an individual limited access to medical care, treatment options and follow up supportive care.

Current clinical management of TBI is primarily determined by the severity of the initial mechanical insult, as pharmacological interventions designed to arrest the secondary injury phase remain largely ineffective.^10^ Standard of care protocols for critical cases prioritize physiological stabilization and specialized intensive monitoring to mitigate subsequent tissue degradation.^11^ While ongoing clinical investigations continue to explore novel neuroprotective agents, such strategies are frequently constrained by high resource requirements and a diagnostic focus on symptomatic relief over direct molecular restoration.^12^ Consequently, there is a compelling requirement for scalable, cost-efficient interventions capable of immediate implementation to address the deleterious, post-traumatic biochemical cascades following trauma.^13^

Nutritional interventions, particularly intermittent fasting (IF), offer a scalable and promising approach to neuroprotection. IF has been investigated for decades across a diverse range of clinical contexts, with foundational research establishing its efficacy in managing obesity, cardiovascular health, and metabolic disorders such as Type 2 diabetes.^14^ Recently, these metabolic insights have transitioned into the TBI landscape.^13^ Human clinical evidence is beginning to confirm the efficacy of metabolic switching in maintaining neural health; for example, current studies are examining whether IF can reduce the levels of neural injury biomarkers, such as neurofilament light (NfL) and glial fibrillary acidic protein (GFAP), following sub-concussive head impacts in athletes.^15^ In human subjects, IF has been shown to improve insulin signaling and decrease systemic inflammation, suggesting it may counteract the metabolic conditions that accelerate neurodegeneration.^16^ The therapeutic value of IF stems from its capacity to function as a metabolic switch that activates endogenous cellular repair pathways, providing a window to either prime the brain for resilience prior to an injury or support recovery in the post-injury period.^16^

Because of the complex pathology associated with TBI, *Drosophila melanogaster* has been used by several groups as a highly robust and high-throughput model system tp study the underlying cellular mechanisms of injury.^17-19^ Our group has developed a traumatic brain injury model using adult *Drosophila*. By utilizing the tissue homogenizer BeadRuptor system, we have produced standardized, closed-trauma conditions that can be applied to adult fruit flies across different ages, genotypes, and therapeutic treatment regimens.^19-22^ In general, we have found that conditions promoting autophagic function, the critical protein aggregate and organelle clearance pathway, in the adult fly nervous system are highly beneficial, leading to trauma resistance and the preservation of neurological function.^20-22^

Other work from our also demonstrated the powerful effect of intermittent fasting (**IF**) treatment on the health-span and stress resistance of *Drosophila*.^23^ Research establishes that IF extends lifespan and delays age-related decline by improving autophagic function in the aging brain.^23^ We previously observed that flies exposed to three weekly 8-hour fasts are significantly protected; on average, wild-type flies and mutant genotypes live longer, demonstrate a reduction in age-related neural aggregates, and maintain baseline performance against the natural decline of adult behaviors.^23^ Neural transcriptomics studies also identified several dynamic expression changes associated with IF treatment, alongside the suppression of transcriptional drift variance in the aged nervous system.^24^ This metabolic mechanism directly aligns with our established TBI framework, indicating that the threshold for post-traumatic mortality is critically linked to the same molecular pathways that govern longevity and the autophagic clearance of protein aggregates.^19,22,23^ By leveraging this TBI platform to reproducibly mirror human phenotypes such as ataxia and progressive neurodegeneration.^18,19^

In this study we have examined the impact that different intermittent fasting treatments have on normal adult flies exposed to traumatic brain injury. We found that pre- and post-IF treatments were protective, promoting longevity and reducing acute and long-term mortality indexes (**MI**). Neural function was also improved, as shown by the suppression of trauma-related defects to climbing behaviors or geotaxis profiles. Using the same IF treatment protocols similar IF-positive trauma responses were also found for aged fly cohorts (3-weeks). Examining the expression of key fly inflammatory markers (AMPs) at critical post-trauma timepoints showed neural levels were suppressed in young (10-day) and aged (21-day) cohorts exposed to either IF-treatment protocol. This suggests the neural protective effect of IF exposure may be linked to suppressing additional secondary damage caused by excessive or protracted inflammation and has wider treatment implications for human head trauma patients.

## Materials and Methods

### *Drosophila* Stocks and Generation of Experimental Flies

The Canton-S and w^1118^ stock lines were originally obtained from the Bloomington Drosophila Stock Center (Bloomington, Indiana), isogenized and maintained as individual stock lines.^19,22,23^ Experimental flies were generated by crossing virgin Canton-S female flies with *w*^*1118*^ males. The resulting F1 male offspring were collected using CO_2_ within four hours of eclosion. Individual cohorts were maintained 30 flies per vial on standard fly media (molasses, agar, cornmeal, yeast) at 24°C with 40-50% humidity on a 12-hour light-dark cycle.^19,22,23^

### Intermittent Fasting Protocol

During IF-treatment studies, flies were maintained using standard conditions and media until exposure to an intermittent fasting regimen using fasting vials (2% agar w/v) for 8 hours per day (0900-1700), 3 days per week on an alternating-day basis excluding weekends.^23^ Both 10-day old and 3-week-old pre-treatment cohorts began the fasting regimen 3 days after eclosion.^23^ Both 10-day old and 3-week-old post-treatment cohorts began fasting 48 hours after 10x mTBI. For all studies, IF-treated flies were placed on *ad libitum* conditions for 24 hours prior to being used for negative geotaxis response assay or being flash frozen in liquid nitrogen and stored at -80°C for subsequent tissue isolation. ^20,23,25^

### Traumatic Brain Injury

Adult fly cohorts were anesthetized using CO_2_, 10 flies were loaded into 2ml screw cap tubes (Olympus Plastics) and allowed to recover prior to being placed into the Omni Bead Ruptor-24 (Omni International, Kennesaw, GA) for traumatic brain injury. Pre-treatment cohort adult flies were aged out to 10-days or 3-weeks while adhering to IF protocol before sustaining 10 mild trauma bouts (10x mTBI, 2.7 m/s).^19,20^ Cohorts of post-treated adult flies were aged to 10 days and 3 weeks on standard food before sustaining trauma (10x mTBI, 2.7 m/s). All injured flies were returned to standard media (30 flies/vial) and cultured for 48 hours before beginning exposed to or returning to intermittent fasting 3x weekly. A set of control (no TBI, no IF) and matching TBI exposed fly cohorts (no IF) were included for each study.

### Aging and Negative Geotaxis Response

Lifespan of each fly cohort (∼100 flies) was recorded by counting the number of dead flies starting 24 hours after 10x mTBI and continuing every 48 hours. Both age cohorts were maintained on standardized intermittent fasting or *ad libitum* conditions alongside age matched controls. The *Drosophila* Negative Geotaxis Response (NGR) relies on the innate escape response of *Drosophila* and uses the Rapid Iterative Negative Geotaxis Apparatus.^19,20^ Groups of 10-30 flies were tapped down, video recordings made and still images taken at 5 seconds. At this timepoint, in climbing index score was counted climbed centimeters, using the ImageJ Cell Counter Plugin.^19,20,23^

### Quantitative Polymerase Chain Reaction

For pre-treatment cohorts, newly eclosed flies were aged to 10 days or 3 weeks on standard fly media before being anesthetized and treated as control (no TBI) or placed on IF protocol prior to being exposed to mTBI conditions and frozen. TBI cohorts were collected 4 h and 24 h after trauma, flash-frozen, and stored (80°C).^19,20,24^ Post-treatment cohorts were aged on standard food prior to mTBI and placed on IF protocol thereafter. Post-treatment flies were allowed to recover for 24 hours on standard food and the placed on either standard food or started on the IF protocol for 3 bouts before being flash-frozen and stored (80°C). Whole heads were isolated, RNA was extracted (TRIzol; Thermo Fisher Scientific,Grand Island, NY), and cDNA libraries were produced (Thermo Scientific, Pittsburg, PA). Quantitative polymerase chain reaction (qPCR) was performed on a CFX Connect Real-Time PCR Detection System (Bio-Rad) using Universal SYBR Mix reagents (Bio-Rad). Primer Sequences for Actin 5c, DptB, and Attc were generated using nBLAST and ordered at 100 μM stocks (Integrated DNA Technologies, Coralville, IA, USA). The Pfalff method was used to quantitate Cq values, load correction, and expression levels. Control (no TBI) mRNA levels were set to 1.0, where trauma cohorts in between and across genotypes were normalized to calculate expression profiles, standard error, and p-values.^19,20,24^

### Statistical Analysis

Behavioral and Longevity profiles, and real time-qPCR were generated using Microsoft Excel, with Figures and Tables assembled in PowerPoint (Mac 2022 v16.69; Microsoft Office). Statistical analyses for longevity, geotaxis, expression profiles, and Western blots were generated using customized R programs. For comparative changes to longevity profiles, p-values were generated with the log-rank test.^20,21^ To compare different NGR cohort profiles, a nested random effects model was used to generate corrected average climbing index, SE, and p-values. P-values for 24-h, 1-w, and 2-w mortality were generated with the Fisher’s exact test, with pairwise p-values adjusted with the Bonferroni correction.^20^ For RT-qPCR data analyses, Kolmogorov–Smirnov and Levene’s tests were performed to identify the appropriate statistical method for mean comparisons.^20^ All normally distributed data were examined using one-way analysis of variances with Fisher’s least significant difference or Games-Howell post hoc tests. For nonparametric data, Kruskal–Wallis tests with Dunn–Bonferroni post hoc tests were used to compare conditions.^20,21^

## Results

We have previously treated traumatized fly cohorts with several different therapeutic compounds, which enhanced average longevity profiles and improved trauma responses (REF). Therefore, we examined the impact that established intermittent fasting and mTBI protocols have on adult *Drosophila* physiology. Fly cohorts (10-days) were exposed to **1)** control, **2)** mTBI (10x), **3)** Pre-IF (mTBI) and **4)** Post-IF (mTBI) conditions, then were maintained on their corresponding treatment platforms for the duration of the study. Flies were fasted 3-times per week, returned to fresh food, with dead flies counted for the remainder of the study. Average group longevity profiles are illustrated as Kaplan-Meier curves, showing both normal and trauma related longevity profiles (**Figure 1A, Table 1A**). Young (10-day) traumatized cohorts that were exposed to Pre-IF and Post-IF treatment showed a partial rescue of the trauma-related phenotypes (REF). Further analysis found that Pre-IF treatment is the most protective (**Figure 1B, Table 1A**). Relative trauma sensitivity is also illustrated by different Mortality Indexes (**MI**) timepoints (24-hr, 48-hr, 1-wk, 2-wk, 3-wk, 4-wk) for each cohort. Pre-IF treated cohorts demonstrate significantly lower MI levels at the 4-week-old timepoint, while Post-IF flies showed similar but not significant mortality trends (**Figure 1C**).

**Table 1.** Young adults (10-days)

| 10-Day Males | Control (TBI 0) | Control 10x mTBI | Pre-IF 10x mTBI | Post-IF 10x mTBI |
| --- | --- | --- | --- | --- |
| mean | 38.64 | 29.20 | 33.69 | 31.92 |
| stdev | 14.99 | 11.83 | 13.34 | 12.21 |
| n | 111 | 106 | 81 | 98 |
| SEM | 1.40 | 1.15 | 1.48 | 1.23 |
| p-val Std Food | - | 7.0546E-08 | 1.1838E-03 | 1.1836E-05 |
| P-val Std + TBI 10x | - | - | 0.0136 | 0.0416 |
| % Change |  |  | +15.4 | +10 |

**Figure 1.**
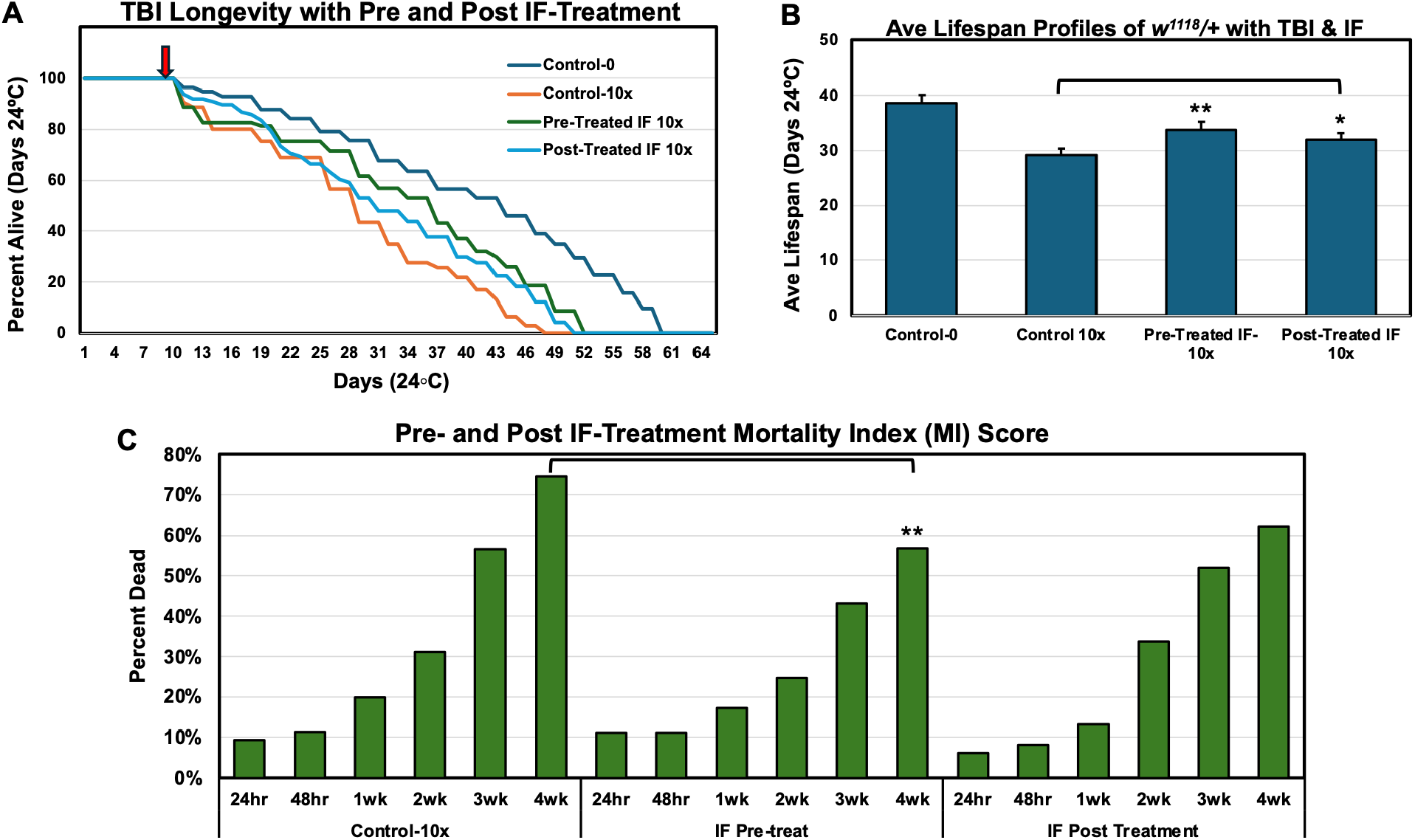
Trauma Longevity and Mortality Profiles with Intermittent Fasting. (a) Kaplan-Meier survival profiles of male wild-type flies (10 days of age) subjected to 10 mild trauma bouts (10x mTBI, 2.7 m/s) with control diet, pre-treated IF, and post-treated IF. Control diet without trauma indicated as Control-0. (b, c) Summary of lifespan profiles for each fly cohort. P-value with respect to control without trauma and with trauma. (d) Mortality index (percent dead) at various time points following trauma. P values are noted as * 0.05, ** 0.01, *** 0.001.

Climbing behaviors were also assessed during aging studies for each fly group, using an established the Negative Geotaxis Response (**NGR**) assay. After trauma exposure, the climbing responses of each fly group were examined at 1 and 2-weeks. Control and traumatized groups showed normal TBI and age-related decline in NGR behaviors (**Figure 2**). While the trauma-related decline wasn’t fully rescued, both IF treatments improved the locomotor behaviors of adult flies (**Figure 2**). Unexcitedly, Post-IF treated flies also showed significant NGR improvement at both the 1 and 2-week timepoints following trauma (**Figure 2**).

**Figure 2.**
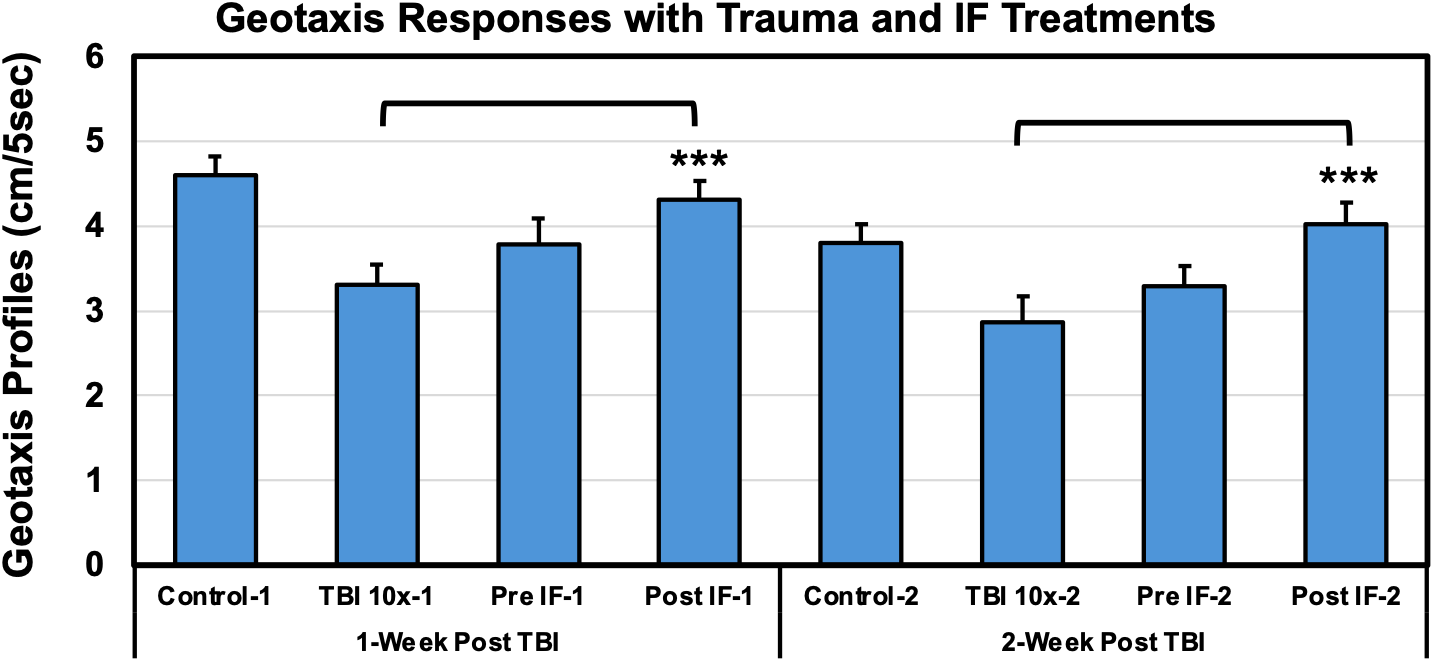
The Negative Geotaxis Response profiles (climbing) of control, TBI and IF-treated fly cohorts at 1-and 2-week timepoints following trauma. P values are noted as * 0.05, ** 0.01, *** 0.001.

Elderly human patients and those with metabolic dysregulation often show complications following traumatic brain injury.^2^ Indeed, we have also found middle-aged flies (3-4 weeks) generally match human responses by having a heightened sensitivity to trauma.^19^ To determine whether intermittent fasting was protective for older adult flies, 21-day-old cohorts were initially tested with less severe mTBI (x5) conditions.^19^ In this case, both IF-treated fly cohorts showed responses that were like those of younger flies (**Figure 3A-D**). Indeed, Pre-IF treatment improved longevity to such an extent that injured flies (21-days) lived longer than matching non-traumatized and mTBI (5x) controls (**Figure 3A-B**). Mortality indexes were significantly improved for Pre-IF flies (**Figure 3C**) and modest improvements at 1-week following a mTBI 5x exposure (**Figure 3D**).

**Figure 3.**
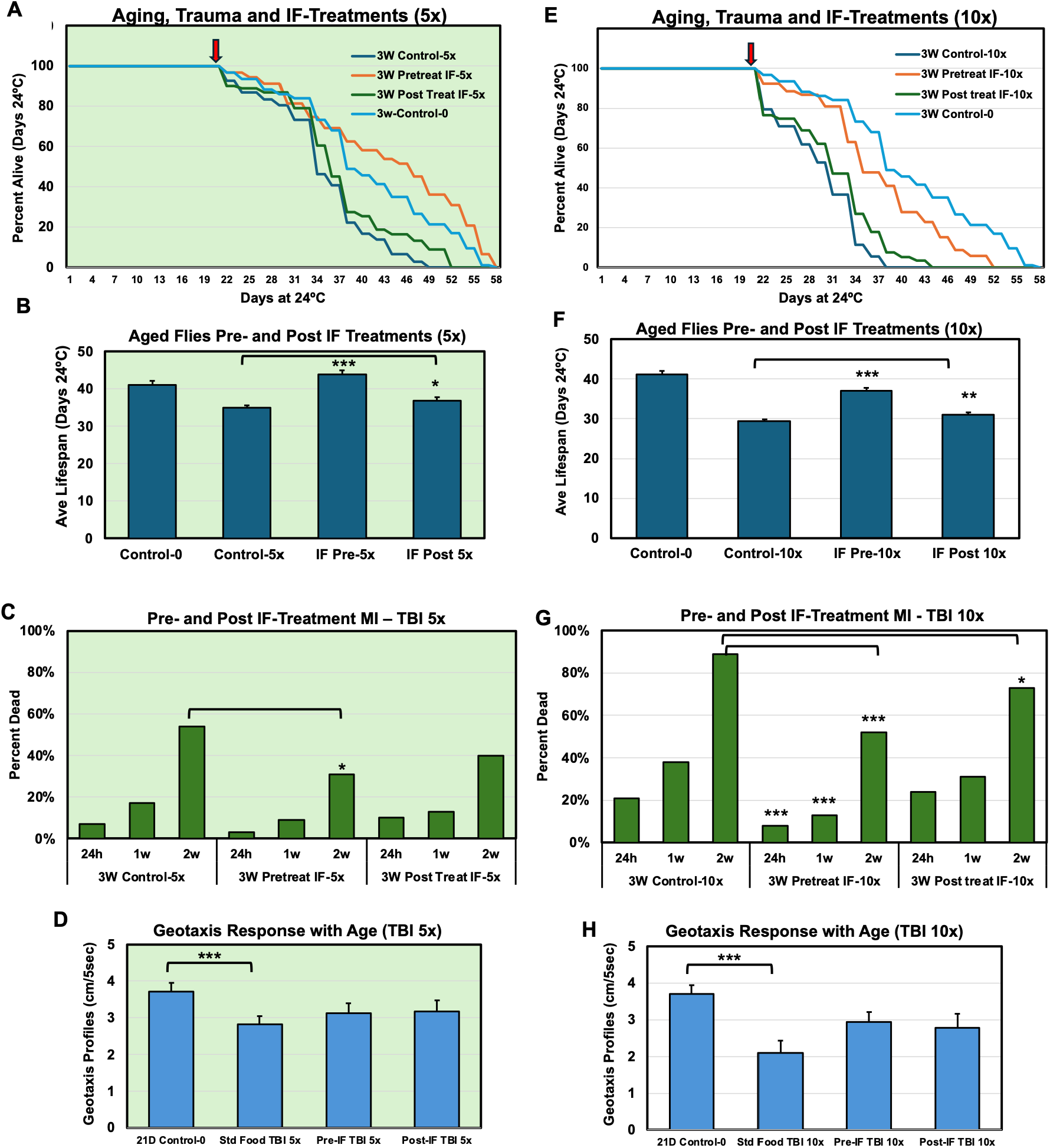
5x or 10x Trauma Bouts and IF-Treatment of Aged Drosophila Males. **A-D)** The profiles for aged male flies (3 weeks of age) exposed to 5 mild trauma bouts (5x) or **E-G**) 10 mild trauma bouts (10x). **A, D**) Kaplan-Meier survival profiles following trauma for control diets, pre-treated IF, and post-treated IF. Controls on a standard diet without trauma indicated as Control-0. **B, E**) Longevity profiles for each fly cohort. P-value with respect to Control 10x mTBI. **C, F**) Mortality index at various time points following IF-treatments and trauma. **D, G**) The trauma related geotaxis responses (cm / 5 sec) of aged flies from each treatment cohort. P values are noted as * 0.05, ** 0.01, *** 0.001.

Since aged flies showed significant improvement, we also examined IF-treatment together with the standard mTBI 10x protocol. Traumatized flies given Pre-IF treatment showed a net positive impact to longevity profiles (**Figure 3D-E**), mortality indexes (**Figure 3F**) and climbing responses (**Figure 3G**). Post-IF treatment, while not as robust, also gave aged flies a measurable positive impact on longevity and behavior profiles (**Figure 3D-F**). Between 21 to 28-days of age is the time when autophagic capacity in the adult *Drosophila* nervous system typically starts to decline and when middle-aged adult flies show increased sensitivity to a variety of stressors. The progressive decrease in neural protein clearance capacity and stress responses is a normal feature of neural aging seen across a range of species, including humans. This indicates that the ability of IF-based diets, to promote longevity and trauma profiles, maybe a conserved and translatable response.^13,14^

**Table 2.** Longevity Profiles of Aged Flies Exposed to Different IF-Treatments and Trauma.

| 21-Day Males | Control (TBI 0) | Control. 10x mTBI | Pre-Treat IF 10x mTBI | Post-Treat IF 10x mTBI | Control 5x mTBI | Pre-Treat IF 5x mTBI | Post-Treat IF 5x mTBI |
| --- | --- | --- | --- | --- | --- | --- | --- |
| mean | 41.09 | 29.44 | 37.01 | 31.02 | 34.98 | 43.80 | 36.88 |
| stdev | 9.63 | 5.18 | 7.93 | 6.34 | 6.76 | 10.92 | 8.01 |
| n | 94 | 131 | 105 | 119 | 108 | 91 | 91 |
| SEM | 0.993 | 0.453 | 0.774 | 0.582 | 0.651 | 1.145 | 0.839 |
| P Values |  |  | 4.82748E-17 | 0.00225 | 1.31476E-08 | 1.23756E-12 | 0.0212 |
| % Change |  |  | +25.7 | +5.3 |  | +25.2 | +5.4 |

Since the adult responses to different mTBI bouts were demonstrated and we were interested in defining the minimal level of intermittent fasting needed to have positive impacts on trauma responses. Aged adult male flies (21-days) were exposed to the standard mTBI, 10 conditions and Pre-IF treated with 0, 1x, 2x or 3x weekly bouts of intermittent fasting. Even a single weekly fast significantly improved longevity (**Figure 4A-B**). An increase or IF dose response was detected, with 2x and 3x weekly IF-treatments resulting greater improvements for each of the trauma assessments tested (**Figure 4A-C**). The implications are that even modest dietary change could have a significant positive impact on trauma related outcomes.

**Figure 4.**
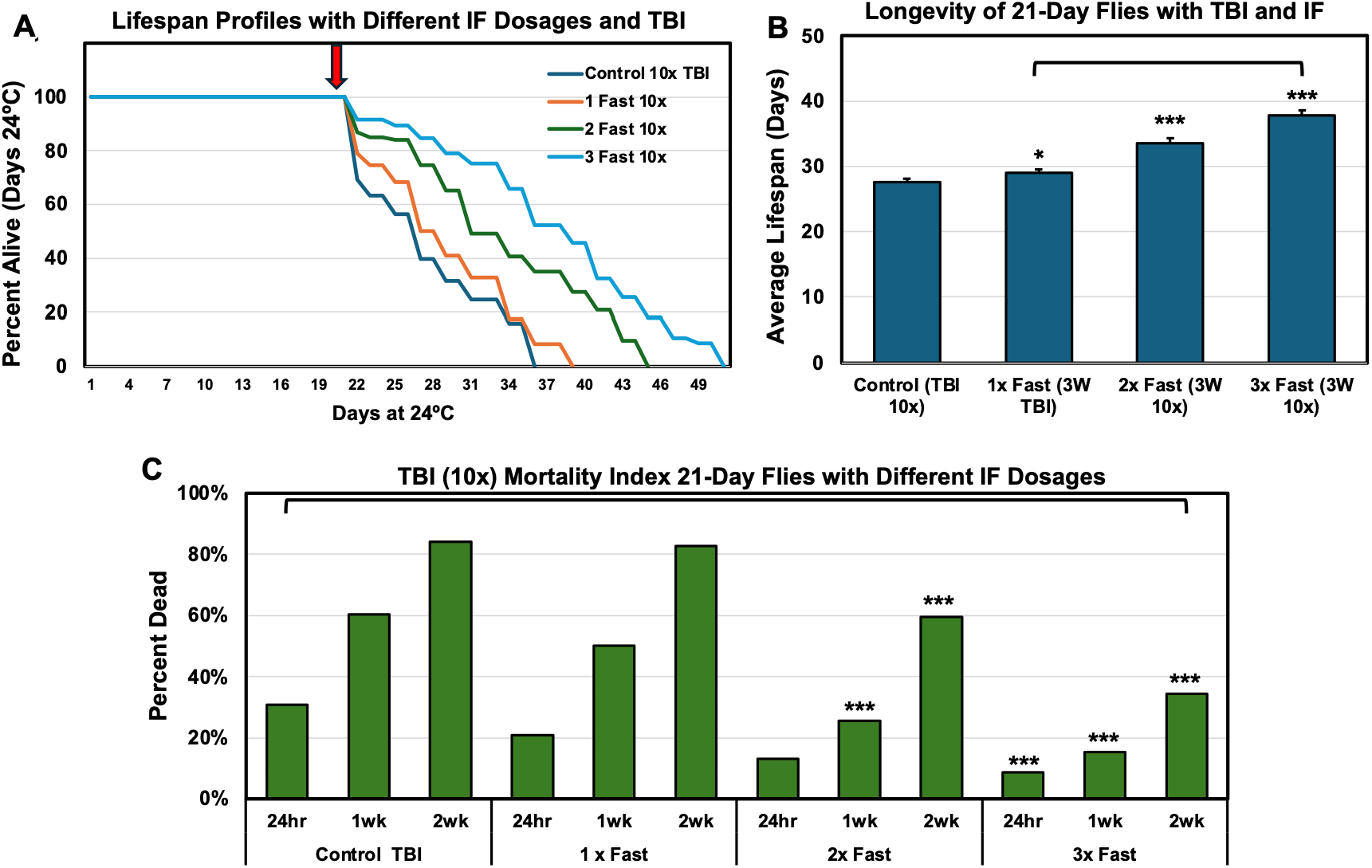
Traumatized Adult Flies Pre-Treated with Different Intermittent Fasting bouts. **A)** Kaplan-Meier curves, **B)** average longevity profiles, **C)** Mortality Index. P values are noted as * 0.05, ** 0.01, *** 0.001.

**Table 3.**
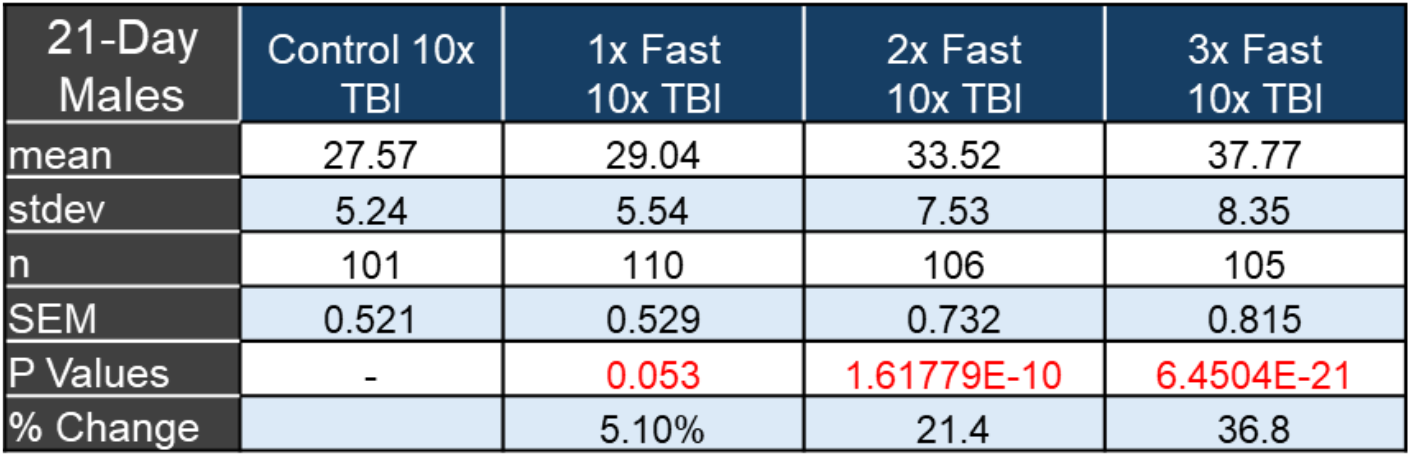
Longevity Profiles of Aged Flies Exposed to Different IF-Treatment Bouts.

| 21-Day Males | Control 10x TBI | 1x Fast 10x TBI | 2x Fast 10x TBI | 3x Fast 10x TBI |
| --- | --- | --- | --- | --- |
| mean | 27.57 | 29.04 | 33.52 | 37.77 |
| stdev | 5.24 | 5.54 | 7.53 | 8.35 |
| n | 101 | 110 | 106 | 105 |
| SEM | 0.521 | 0.529 | 0.732 | 0.815 |
| P Values | - | 0.053 | 1.61779E-10 | 6.4504E-21 |
| % Change |  | 5.10% | 21.4 | 36.8 |

A feature often seen with human trauma patients is a marked increase in several inflammatory markers, including elevated levels of several (IL-6, IL-1β), TNF-α and C-reactive protein (CRP).^26-28^ Our group and others have found several *Drosophila* antimicrobial peptides (AMPs), which are highly responsive components of the innate immune systems, behave as inflammatory markers following stressors like TBI.^19,20^ In the nervous system several AMP genes have a highly robust and choreographed timeline of expression changes following trauma.

Typically, AMP levels start to increase 1-hr following trauma, peak at 4-hrs and typically decline by the 24-hrs post-TBI timepoint (10x).^19^ Here *ad libitum* controls showed *AttC* and *DptB* message levels increased significantly at 4-hr Post-TBI timepoint (**Figure 5A**). This was followed by *AttC* expression falling 24-hrs following mTBI (**Figure 5A**).

**Figure 5.**
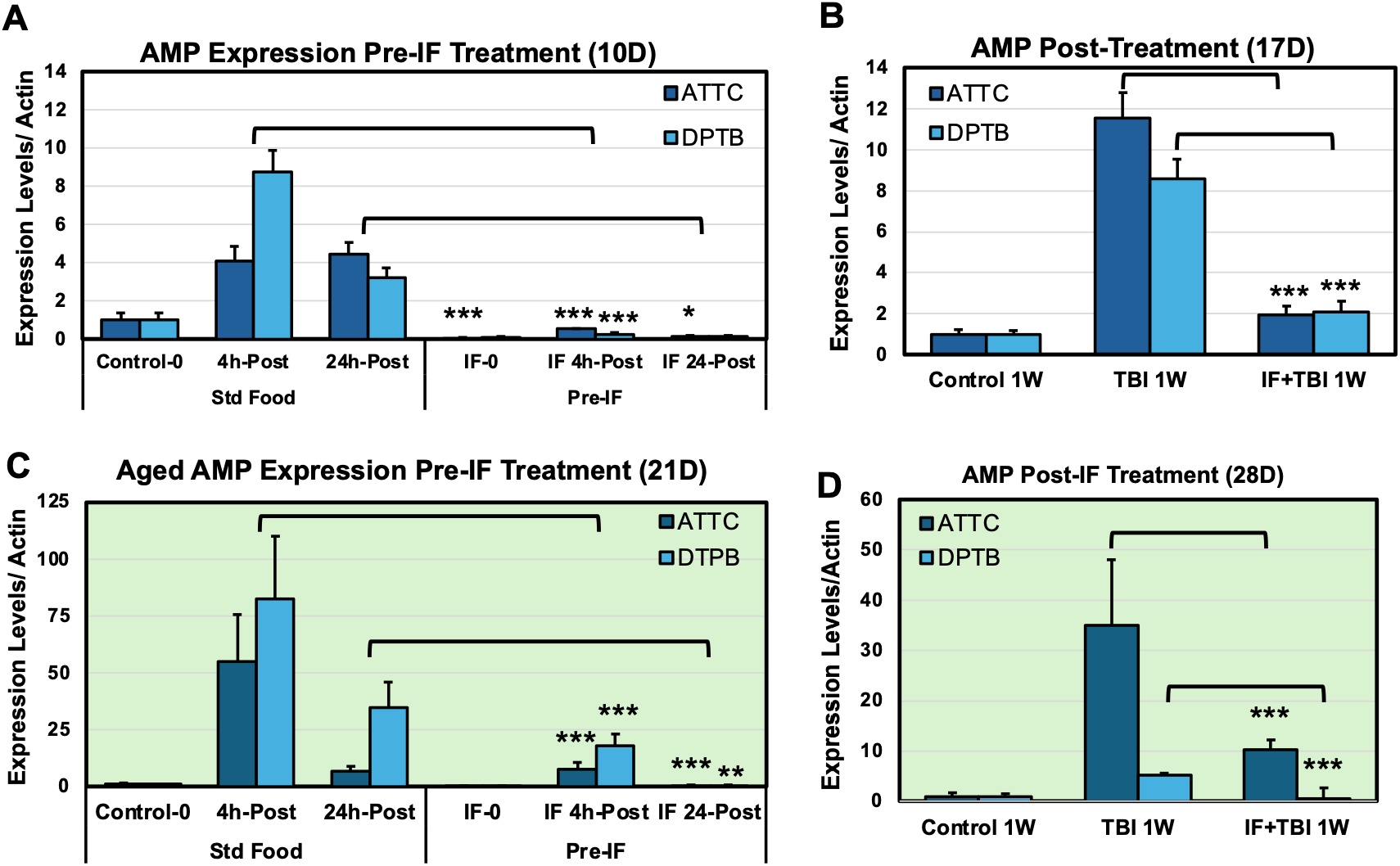
AMP Expression Profiles Following mTBI (10x) Exposure and Pre- and Post-IF Treatments. Gene expression profile of AMP inflammatory markers, AttC and DptB. AMP levels are normalized to 1 at the 0hr timepoint. **A**) AMP levels for young (10D) pre-treated IF flies examining 0hr, 4hr, and 24hr following trauma. **B**) AMP levels for post-treated IF flies at 1 week following trauma. **C-D**) AMP expression profiles of pre-and post-IF treated AMP profiles in aged Drosophila cohorts (21D). P values are noted as * 0.05, ** 0.01, *** 0.001.

The expression profiles of young Pre-IF treated fly cohorts (IF-0, 10-day) were examined and started with a significantly lower baseline level of *AttC* in neural tissues (**Figure 5A**). After trauma exposure flies had significant attenuation of AMP message production at the IF-4-hr and IF-24-hr timepoints (**Figure 5A**). To examine Post-IF impact on AMP expression profiles, young flies were also examined 1-week following trauma (17-day). Compared to matched baseline controls, 1-week following TBI produced a significant increase in AMP expression, while Post-IF treated cohorts had significantly lower neural AMP profiles (**Figure 5B**). For aged flies (21-day) there was marked AMP elevation, together with increased variability at the IF-4-hr and IF-24-hr timepoints following trauma (**Figure 5C**). Aged IF-Post treated flies again showed lowered neural AMP mRNA levels 1-week following injury (**Figure 5C**). Overall, the positive impact on longevity and behaviors by both intermittent fasting condition is consistent with the neural protection that is afforded by lowering age and trauma related inflammation. This suggests the positive trauma related impact for the recovering CNS maybe that IF protects neurons from secondary rounds of damage due to excessive inflammation.^20,22^

Marcus

## Discussion

### A. Human Trauma Responses, Age, and Metabolic Vulnerability

Traumatic brain injury (TBI) remains a major neurological cause of morbidity and mortality amongst various human populations. Clinical results following an initial mechanical trauma are substantially worsened by an extended secondary phase of injury, that is defined by ongoing neural attrition, systemic metabolic issues, and a failure of proteostatic mechanisms.^12^ This post-traumatic susceptibility is notably evident within an elderly demographic. Reports from the CDC indicate that older people, aged 75 and older account for a higher percentage of TBI-related hospitalizations (32%) and deaths (28%), with unintentional falls being the primary driver of these statistics.^2,3^ It is important to note that older populations possess reduced baseline cellular and neural resilience and a modified metabolic landscape, leaving the aged nervous system uniquely vulnerable to progressive neural degeneration over time, even after a relatively minor or repetitive series of mTBI events.^4^

In addition to chronological age, metabolic factors such as obesity, Type 2 diabetes, and long-term consumption of diets high in sugar and saturated fats serve as major drivers of rapid clinical decline.^29,30^ These dietary habits disrupt mitochondrial function, impair insulin signaling, and prime the innate immune response for over-activation.^16^ Consequently, when a metabolically compromised or elderly brain sustains trauma, secondary injury responses are greatly increased.^12^ While sufficient caloric intake and balanced nutrition are necessary immediately following injury for wound healing, chronic metabolic syndrome instead promotes persistent neuroinflammation. This metabolic priming of inflammatory responses prevents proper tissue recovery and speeds up long-term neurological decay, highlighting the requirement for scalable interventions to address how diet governs head trauma outcomes.^13^

### B. The Drosophila Model in Neuroprotective Discovery

To systematically investigate the molecular pathways connecting diet, neural aging, and trauma, *Drosophila melanogaster* provides a robust and high-throughput genetic model.^17,19^ The conservation of metabolic factors and innate immune defenses between fruit flies and humans makes this platform well-suited for translational research. Utilizing the tissue homogenizer BeadRuptor system to produce standardized, closed-trauma conditions allows us to have precisely control over a range of injury parameters that includes substantial number of individual within a cohort that represents combinations of various treatments, ages and/or genotypes. This methodology reproducibly mirrors human phenotypes such as ataxia, and the progressive decline of other behaviors including sleep disruption.^19^

Previous work from our group examining neural protective measures has focused on pharmacological agents to reduce post-traumatic deficits. A common theme from these studies is that interventions that maintain neurological function with autophagic function. Our work previously demonstrated that the immediate post-injury survival threshold, measured as the mortality index at 24 hours, is linked to the same molecular machinery that regulates health span and longevity.^19^ Because intermittent fasting acts as a powerful metabolic switch capable of maintaining neural autophagy and suppressing transcriptional drift in the aged nervous system, examining IF regimens became our next focus.^24^ By evaluating timed fasting windows, we sought to drive endogenous neuroprotection using a framework that avoids pharmacological intervention.^20-22^

### C. Prophylactic and Therapeutic Efficacy of Intermittent Fasting Across Age Cohorts

Our findings across both young (10-day-old) and aged (3-week-old) fly cohorts provide evidence that intermittent fasting acts as both a protective priming mechanism and a post-injury restorative treatment. Flies pre-treated with IF before sustaining trauma, exhibited improved trauma resistance and lowered acute mortality indices (MI).^20^ Crucially, starting the IF protocol after sustaining injury also suppressed the secondary injury phase, helping to maintain neurological function and increasing lifespan compared to *ad libitum* standard food controls. The preservation of neural health is most striking in our functional assessment of climbing behaviors, specifically the negative geotaxis response profiles. While standard diet flies showed a rapid, trauma-related decline in coordination, both Pre- and Post-IF treated cohorts demonstrated robust suppression of behavioral deficits regardless of age.^19,20,25^ To optimize this effect, we mapped out dose-response profiles by varying fasting frequency. Our data shows that neuroprotective efficacy depends on reaching a metabolic threshold to activate systemic pathways, with optimized protocols reducing mortality.

### D. Neural Inflammatory Regulation with Age and Trauma

We found that inflammatory signaling was significantly reduced for both Pre- and Post-IF groups in both the young and aged fly cohorts. As shown by their expression profiles, the aged fly cohorts demonstrated an increase in inflammatory signaling accompanied with reduced neurological function.^19^ The progressive decrease in neural protein clearance capacity and autophagy is a normal feature of neural aging seen across a range of species, including humans.^25,31^ The ability of restrictive diets, including intermittent fasting to improve longevity in flies, parallels findings in human populations.^32^ Given that elderly individuals are disproportionately vulnerable to poor TBI recovery, these findings imply the usage of IF as a scalable, inexpensive intervention for improving trauma resilience preventatively and restoratively.

## Conclusions

In *Drosophila* model, intermittent fasting demonstrates a significant improvement to adult longevity and behavioral outcomes while reducing inflammatory signaling when compared to control diet. Our study investigated this finding further, demonstrating its scalable efficacy with quantity of fasting intervals as well as its effectiveness amongst aged fly cohorts. With elderly human populations facing higher risks related to trauma, IF can offer an accessible solution to providing greater outcomes. Clinical trials experimenting with IF in human populations will be greatly beneficial to creating a defined protocol to help such populations at risk. Additionally, future studies should aim to understand the causal link between intermittent fasting reduced inflammatory signaling such as through a genetic knockdown.

## Notes

### Competing Interest Statement

The authors have declared no competing interest.

